# Bromodomain protein inhibition protects β-cells from cytokine-induced death and dysfunction via antagonism of NF-κB pathway

**DOI:** 10.1101/2020.11.05.363408

**Authors:** Vinny Negi, Jeongkyung Lee, Ruya Liu, Eliana M. Perez-Garcia, Feng Li, Rajaganapati Jagannathan, Ping Yang, Rita Bottino, Ke Ma, Mousumi Moulik, Vijay K Yechoor

## Abstract

Cytokine induced β-cell apoptosis is the major pathogenic mechanism in type 1 diabetes (T1D). Despite significant advances in understanding underlying mechanisms, few drugs have been translated to protect β-cells in T1D. Epigenetic modulators such as bromodomain-containing BET (Bromo- and Extra-Terminal) proteins are important regulators of immune responses. Pre-clinical studies have demonstrated a protective effect of BET inhibitors in NOD (non-obese diabetes) mouse model of T1D. However, the role of BET proteins in β-cell function in response to cytokines is unknown. Here we demonstrate that I-BET, a BET protein inhibitor, protected β-cells from cytokine induced dysfunction and death. In vivo administration of I-BET to mice exposed to low-dose STZ (streptozotocin), a model of T1D, significantly reduced β-cell apoptosis and preserved β-cell mass, suggesting a cytoprotective function of I-BET. Furthermore, human islets treated with I-BET displayed better glucose stimulated insulin secretion compared to controls, when exposed to cytokines. Mechanistically, RNA-Seq analysis revealed I-BET treatment suppressed pathways involved in apoptosis, including NF-kB signaling, while maintaining the expression of genes critical for β-cell function, such as Pdx1 and Ins1. Taken together, this study demonstrates that I-BET is effective in protecting β-cells from cytokine-induced dysfunction and apoptosis, and may have potential therapeutic values in T1D.

## INTRODUCTION

Type 1 diabetes (T1D) is an autoimmune disorder characterized by the destruction of insulin producing β-cells (1–3). At molecular level, inflammatory cytokine leads to its dysfunction with impaired glucose stimulated insulin secretion (GSIS), decreased proliferation, and increased apoptosis. While pancreas or an islet transplant leads to ‘cure’ and insulin-free periods, they remain challenging due to limited donor availability and the need for chronic immunosuppression (4; 5). Hence, insulin replacement remains the standard of care to date, although the difficulty of long-term injections to maintain good glycemic control makes insulin therapy suboptimal. Replacing β-cells with those derived from patient-specific iPS (induced pluripotent stem) cells remains in experimental stages and yet to overcome the hurdle of cytokine induced apoptosis in an autoimmune setting (6; 7). Thus, there is an imperative need for developing therapeutics for T1D that protects β-cells from cytokine-induced cell death to preserve β-cell mass.

The bromodomain-containing BET (Bromodomain and extra-terminal domain containing) proteins, including BRD2, BRD3, BRD4, and BRDT, are epigenetic regulators of gene transcription known to be involved in inflammatory response, proliferation, apoptosis, and differentiation (8; 9). These proteins bind to the acetylated lysine of histones in the genome, acting as a ‘reader’ and recruit histone remodeling proteins, such as HATs (histone acetyltransferase), HDACs (histone deacetylase), and transcription elongation factors (9; 10). Recently, with the advent of a new generation of higher affinity chemical inhibitors such as I-BET, there has been a surge in testing their therapeutic utility for many oncological (prostate, breast, hematopoietic glioblastoma) and vascular (pulmonary arterial hypertension) diseases despite the limited knowledge of underlying molecular mechanism (11; 12); I-BET primarily targets BRD2, 3, and 4 (8; 13).. In diabetic NOD mice, I-BET was shown to promote an anti-inflammatory phenotype of infiltrating pancreatic macrophages and enhance β-cells proliferation (14), with a reduced SASP (senescence-associated secretory phenotype) response (15), though a cell-autonomous effect in β-cells was not established. At the cellular level, BET inhibition by JQ1 in INS1 cells led to an increase in insulin content and secretion, though this was assessed only under basal conditions (16). Whether inhibition of BET proteins in β-cells could confer protection to cytokine-induced injury remains unclear. Thus, to understand its potential therapeutic potential in T1D, it is crucial to determine the effect of BET inhibition on protecting β-cell function and identify underlying molecular mechanisms.

In this study, we used I-BET762, from here on referred to as I-BET, as a representative BET protein inhibitor that is currently in clinical trials, to test its protective effects on β-cells (11; 17). Employing both in vitro and in vivo approaches using low dose STZ model of T1D, we specifically determined that I-BET had a protective effect against cytokine-induced apoptosis and β-cell dysfunction. Mechanistically, using global transcriptomic analysis, we determined I-BET treatment not only enhanced critical factors that promoted β-cell function but also antagonized the cytokine-induced upregulation of the NF-kB pathway that is known to induce apoptosis in pancreatic β-cells in T1D (18).

## RESEARCH DESIGN AND METHODS

### Cell culture

Rat insulinoma, INS1 (832/13 cells a gift from Dr. Christopher Newgard) cells were cultured in RPMI-1640 medium containing 11mM glucose, 10% fetal bovine serum, 10mM Hepes, 2mM L-glutamine, 1mM sodium pyruvate, and 0.05mM 2-mercaptoethanol. The cells were pre-treated with I-BET762 (500nM, SelleckChem) or vehicle control (VC) for 48h followed by the addition of cytokine cocktail (CC) of IL-1β (10 ng/ml), IFN-γ (100 ng/ml), and TNF-α (25ng/ml)(Peprotech) for another 8h or 24h as indicated in the presence of I-BET or VC.

### Annexin-PI apoptosis assay

INS1 cells, after treatment as indicated, were trypsinized and stained with Annexin-V (APC labelled) and PI for 15 min at room temperature in dark and assessed on flow analyzer BD LSR II using FACS Diva software and analyzed on FlowJo.

### RNA isolation and real-time PCR

RNA was isolated using Direct-zol RNA miniprep kits from Zymo Research according to manufacturer instructions and quantified using Nanodrop. The cDNA was synthesized using amfiRivert kit from GenDEPOT according to manufacturer instructions. The qRT-PCR was performed using amfiSure qGreen master mix from GenDEPOT according to manufacturer instructions on Quant Studio 3 from Applied Biosystems. All the primers used are listed in **Supplementary Table 1**.

### Glucose stimulated insulin secretion (GSIS) and insulin content determination

GSIS was performed as previously reported (19). Briefly, INS1 cells treated with I-BET and CC as indicated and incubated in KRB (Krebs Ringer Biocarbonate) buffer with no glucose for 30min at 37°C with CC and I-BET/VC, followed by serial incubations for 30 min in 2.8mM, 11.1mM and 25mM glucose in KRB. The supernatant was collected for insulin secretion assay. At the end of the experiment, to determine cellular insulin content, the cells were lysed with acid-ethanol lysis buffer overnight at 4°C and supernatant collected after centrifugation at 16000 rcf for 30 min. Insulin secretion and content were determined with the insulin ELISA kit (Crystalchem) according to the manufacturer instructions and normalized to the DNA content.

### RNA Seq

The integrity and purity of isolated RNA was analyzed using RNA Nano 6000 Assay Kit of the Bioanalyzer 2100 system (Agilent Technologies, CA, USA). A total amount of 1 μg RNA per sample was used as input material for sample preparations. Sequencing libraries were generated using NEBNext®Ultra™RNA Library Prep Kit for Illumina®(NEB, USA) sequenced on an Illumina platform (Novaseq 600) according to manufacturer’s instruction and 125 bp/150 bp paired-end reads were generated. Raw data obtained were filtered, cleaned, and aligned to genome using HISAT2. The gene expression was calculated as FPKM and differentially expressed genes across all the samples with p_adj_<0.05 were represented as a heatmap. The gene overlap between the two groups was determined using Oliveros, J.C. (2007-2015) Venny. Gene set enrichment analysis (GSEA) (20; 21) was performed among CCVC (CC+VC) and CCIB (CC+I-BET). The pathway enrichment was determined by DAVID bioinformatic resources 6.8 (22; 23).

### Western blotting

Western blotting was performed as described in (24). Briefly, lysates were prepared in ice-cold 1X lysis buffer (ThermoFisher Scientific) supplemented with 1X protease inhibitor and phosphatase inhibitor (Roche). An equal amount of protein samples was loaded and fractionated by SDS-PAGE, then transferred to 0.2 μm pore-size nitrocellulose membranes (Millipore). Membranes were blocked in 5% non-fat milk-TBS (w/v), then incubated with indicated primary antibodies-cleaved caspase-3 (CST, 9664), Pdx1 (Abcam, ab47308), phospho-p65, p65, p-IKKA/B, IKKB, p-IκBα, IκBα (CST, 9936), overnight at 4°C, later with DyLight 680 or DyLight 800 - conjugated secondary antibodies. The blots were imaged using Licor Odyssey Clx and quantified by Image Studio ver 5.2 software.

### Mouse studies

All animal experiments were approved by the Institutional Animal Care and Use Committee of the University of Pittsburgh. 8week old C57BL/6N mice were purchased from Jackson Laboratory. For induction of diabetes, 50mg/kg of streptozotocin (STZ), freshly dissolved in Na-citrate buffer at pH4.2, was administered via i.p. for 5 days. The mice were distributed into 3 groups: VC-Non-diabetic controls (Sodium citrate buffer only), STZ+VC – STZ with vehicle control for I-BET, and STZ+I-BET; with n=9 in each group. The drug I-BET or its vehicle control (2% DMSO) was given daily orally by gavage for 4 weeks. The body weight and blood glucose level of all the mice were monitored every week for 4 weeks. Glucose tolerance test (GTT) was performed on day 26 and 27 in overnight fasted mice by administering 1.5g/kg of body weight by i.p. of D-glucose. Mice were euthanized on day 28 and pancreas was collected. The blood glucose was measured using a glucometer and by Infinity glucose hexokinase from ThermoFisher Scientific according to the manufacturer’s instructions.

### Immunofluorescence staining

Mice pancreatic tissue fixed in formaldehyde were embedded in paraffin and sectioned at 5 μm thickness. All the sections were stained as previously reported (24). Briefly, slides were warmed for 20 min in 60°C followed by deparaffinization using citrasolv, then rehydrating with decreasing ethanol concentrations. This was followed by permeabilization in 0.25% of Triton-x-100 in PBST for 5 min and antigen retrieval in sodium citrate buffer pH6 in a microwave (1 min at maximum power, then 20 min at minimum power). After cooling down to room temperature they were blocked with 1% BSA in 1x PBS and incubated with primary antibodies against cleaved caspase-3 (CST-9661, 1:100), INS1 (Abcam-ab6995, 1:400) overnight at 4°C and subsequently labeled with secondary antibodies for 1h at room temperature. All the images were taken by the Nikon Fluorescence microscope.

### β-cell mass determination

Two slides, per mice, at least 50 microns apart were selected to stain for Insulin. Images of whole sections were taken by automated EVOS FL Auto 2 imaging system. For each image insulin-positive area was measured and the number of islets counted manually. The size of the islets was categorized into small, medium, and large based on their arbitrary area as computed by ImageJ - less than 10,000 (small), 10,000-30,000 (medium), and greater than 30,000 (large).

### Human islets

Human islets were provided by IIDP and Allegheny health network institute of cellular therapeutics. The details of the donors are described in **Supplementary Table 2**. Received human islets were collected by centrifugation and cultured in CMRL1066 (Mediatech, Corning) supplemented with 10% human serum (Sigma), 1% P/S and 2 mM L-glutamine. They were pre-treated with I-BET762 (500nM) or vehicle control (VC) for 48h followed by the addition of cytokine cocktail (CC) of IL-1β (10 ng/ml), IFN-γ (100 ng/ml), and TNF-α (25ng/ml) for another 24 h in presence of the I-BET or VC. GSIS was done as described earlier using equal number (5-6) and similar sized islets per group.

### Statistical analysis

The data are presented as means ± sem, the in vitro data is representative of three independent experiments. Significance tests were determined by either one-way or two-way ANOVA followed by either Sidak’s multiple comparisons test. p< 0.05 was considered statistically significant. GraphPad Prism 8.3.1 software was used for statistical analyses.

### Data and resource availability

The RNA-seq raw and processed datasets have been deposited in GEO database (GSE160573).

## RESULTS

### I-BET protects against cytokine inducedβ-cell dysfunction

Cytokine-induced β-cell dysfunction with impaired insulin secretion and eventual cell death is the defining feature of T1D. To mimic this in vitro, INS1cells were exposed to a cytokine cocktail (CC) of IL-1β (10 ng/ml), IFN-γ (100 ng/ml), and TNF-α (25 ng/ml) for 8h to determine effects on gene expression and GSIS, or for 24h to analyze apoptosis, respectively. I-BET or DMSO vehicle control (VC) were administered to INS1 cells for 48h followed by cytokine exposure. Gene expression analysis revealed that I-BET treatment significantly up-regulated expression of β-cell function related genes, including *Ins1*, *Ins2*, *Pdx1*, *MafA*, and *Pax6*. More importantly, it was able to prevent the repression of these genes induced by cytokine exposure (**Figure 1A-E**), suggesting cytoprotective effects of I-BET upon cytokine challenge. Notably, I-BET largely restored the expression of these genes to levels that were comparable without cytokine exposure. The regulation of *Pdx1* mRNA was also reflected in a consistent increase in Pdx1 protein levels with I-BET treatment (**Figure 1F-G**). Interestingly, despite changes in mRNA expression MafA protein levels were not altered by cytokine or I-BET treatments after 8 h cytokine exposure, suggesting that post-translational modifications and stability of MafA protein may impact acute changes from cytokine exposure (25; 26) (**Figure 1H-I**). Interestingly, the expression of other genes such as *Pax4, Nkx2.2*, and *Nkx6.1* did not respond to I-BET pre-treatment (**Supplementary Figure 1A-C),** suggesting potential specificity of gene regulation by BET proteins. We then assessed if these gene regulations induced by I-BET led to functional improvement in GSIS. Under basal (2.8mM glucose) condition, insulin secretion was increased by both I-BET and cytokines (**Figure 1J**) at the early 8h time point, but not at 24h (**Supplementary Figure 2**). More importantly and in concordance with the changes seen in the expression of *Pdx1* and *Ins1*, I-BET prevented the decrease in insulin secretion index induced by cytokine exposure (**Figure 1K**). Total cellular insulin content was increased by pre-treatment with I-BET, as was reported by others (16), but this was lost in the presence of cytokines (**Figure 1L**). However, at 24h of cytokine exposure, I-BET was unable to improve β-cell function as determined by gene expression and GSIS (**Supplementary Figure 2**), a possible reflection of the induction of apoptosis with longer cytokine exposure (described below). These data demonstrate that pretreatment with I-BET protected INS1 cells from the deleterious effects from cytokines at an early timepoint but unable to do so at a later stage.

**Figure 1.**
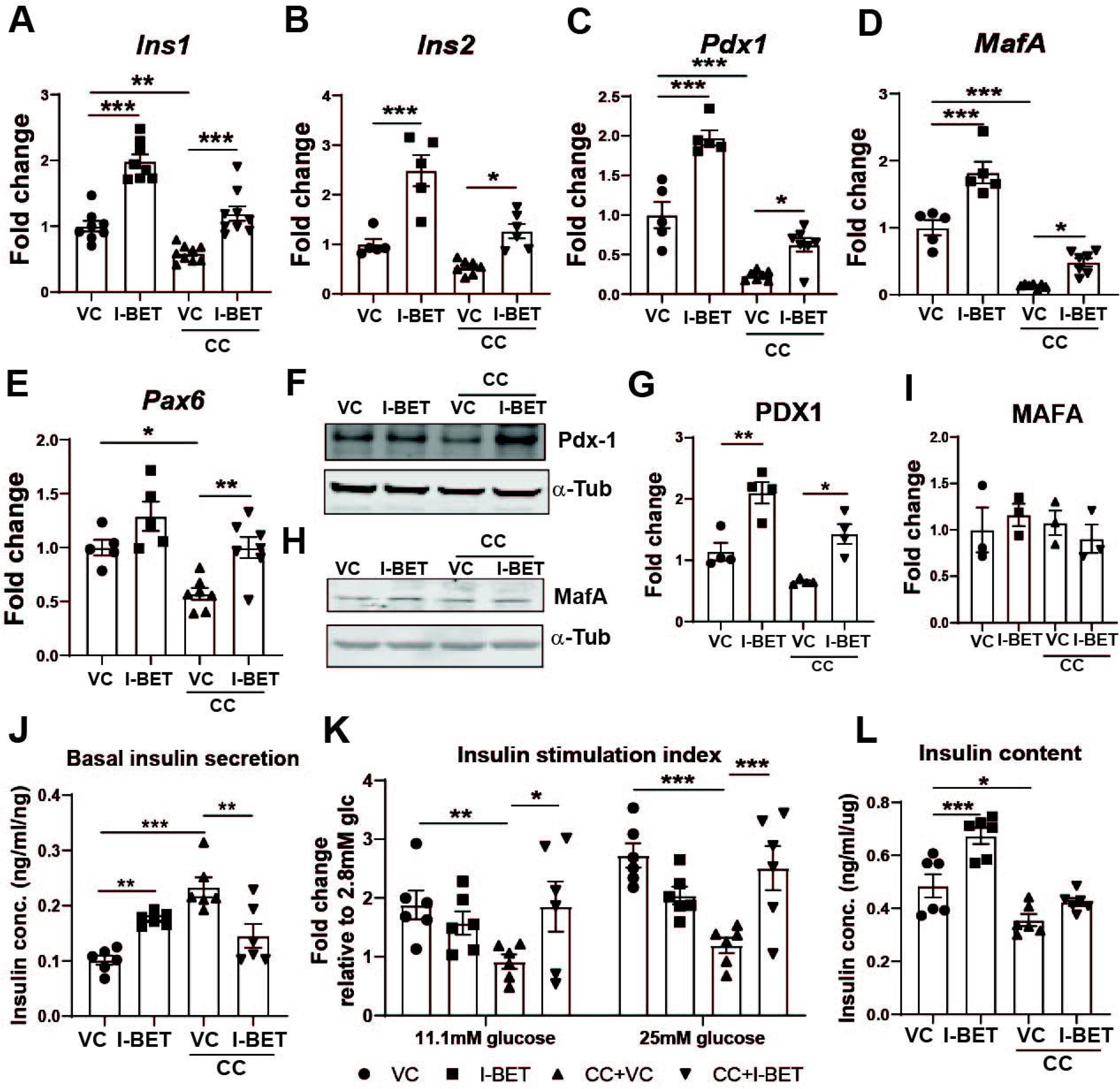
I-BET protects against cytokine induced β-cell dysfunction. INS1 cell were pretreated with I-BET or VC (vehicle control) for 48h and then cytokine cocktail (CC) was added for another 8h. **(A-E)** Gene expression by RT-qPCR of (**A**) *Ins1*, (**B**) *Ins2*, (**C**) *Pdx1*, (**D**) *MafA*, and (**E**) *Pax6* are shown after normalization to housekeeping gene as a fold change over VC. **(F-I)** The protein level by western blotting and the quantification by densitometry of 4 independent experiments is shown for Pdx1 (F, G) and MafA (H, I) and represented as a fold change over VC. α-Tubulin was used as a loading control. **(J-K)** The secreted insulin from INS1 cells in basal 2.8mM glucose (J) and after incubation in indicated glucose concentrations (K) is shown. Insulin secretion is represented as insulin stimulation index (K) a fold change over the respective levels from the basal 2.8mM glucose. (**L**) Insulin content measured in INS1 cell lysates normalized to cellular DNA. The data is represented as mean ± sem (n=3-7) with at least three independent experiments. Statistical significance was calculated using one-way Anova; ***p<0.001, **p<.01, *p<0.05.

### Global transcriptome profile reveals β-cell function related genes were rescued by I-BET treatment

To determine the transcriptional mechanisms underlying I-BET protection of β-cell function in the presence of cytokine exposure, we performed RNA-Seq analysis to identify the enriched pathways regulated by I-BET. Among the four groups - VC (vehicle control), IB (I-BET762), CCVC (cytokine cocktail+ vehicle control), and CCIB (cytokine cocktail+ I-BET762), 9694 genes were differentially expressed (**Figure 2A**). Further analysis revealed that 2932 were up-and 2750 genes down-regulated by cytokines (CC+VC) as compared to VC. 14% of both the up- and down-regulated genes were reverted by I-BET treatment (**Figure 2B**). Interestingly, ER processing and MODY (maturity onset diabetes of the young) related pathways were among the most enriched that were reversed by I-BET treatment (**Figure 2C**). In contrast, metabolic pathways and oxidative phosphorylation were enriched by cytokines exposure but not rescued by I-BET (**Supplementary Figure 3**). Comparison of genes differentially regulated by I-BET with or without cytokines revealed 45.3% overlap of genes, which belong to processes critical for maintaining normal β-cell function and insulin secretion, including ER protein processing, FOXO1 signaling, MODY, and vesicle trafficking (**Figure 2D**).

**Figure 2.**
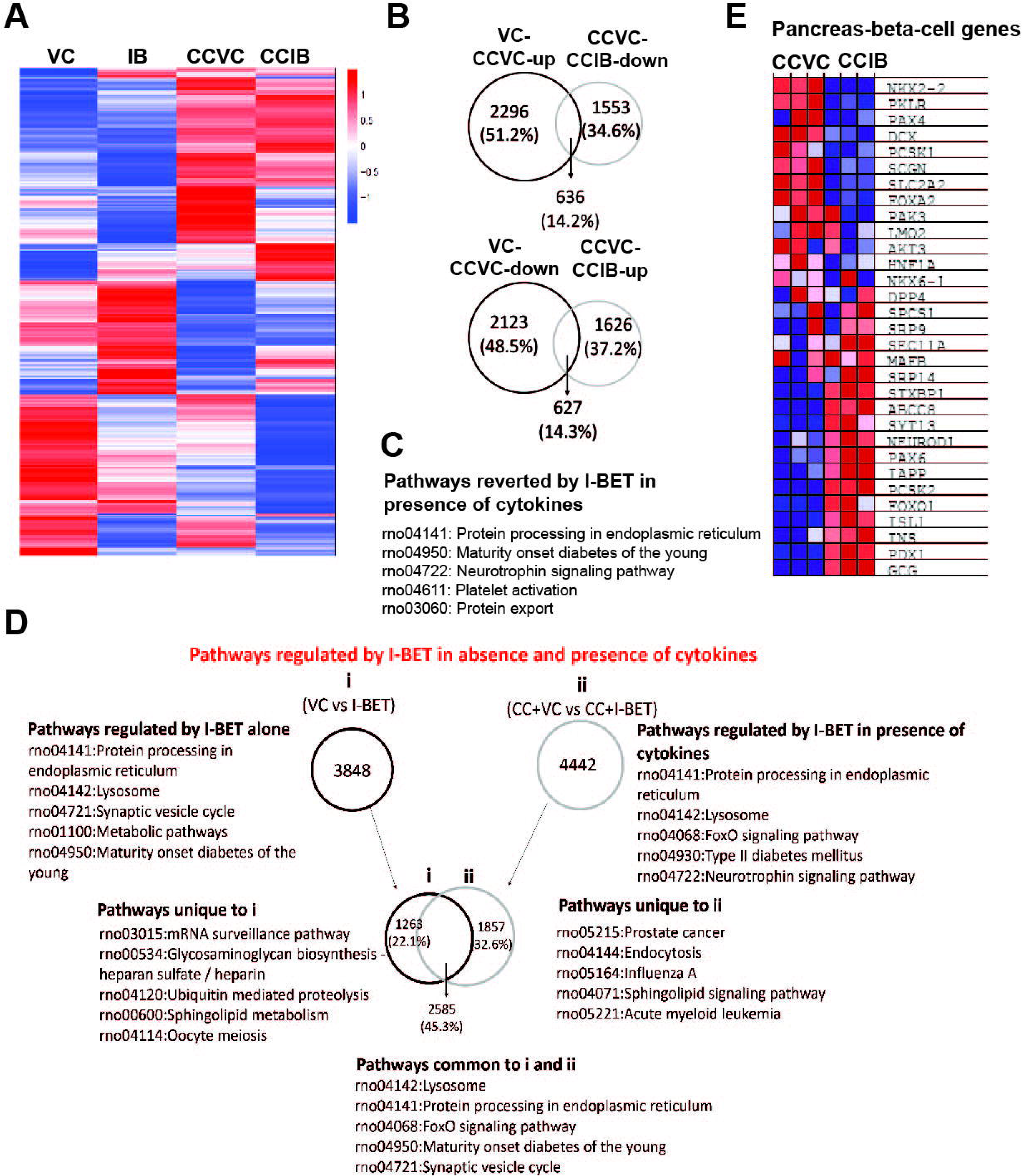
Global transcriptomic changes after I-BET rescue of cytokine-induced changes in INS1 cells. **(A-D)** RNA-Seq analysis from INS1 cells treated with I-BET or vehicle (VC) under short (8h) exposure to cytokine cocktail (CC). (**A**) 9694 genes were found to be differentially expressed among four groups as demonstrated by the heat map. The red and blue colors indicate up- and down-regulated genes expression as measured by log2(ratio) of FPKM. **(B)** Among the genes that were differentially regulated by cytokine cocktail (VC vs CCVC), only 14.2% were reversed by I-BET (CCVC vs CCIB). **(C)** The top 5 enriched pathways for those genes as determined by DAVID included ER processing and maturity onset diabetes of the young (MODY) as most enriched pathways. (**D**) Pathway enrichment by DAVID of genes that were regulated by I-BET alone (IB) and in presence of CC (CCIB). Further with the help of Venn diagrams, they were divided into those that were unique and common to these conditions. **(E)** The heatmap of genes strongly associated with β-cell function are represented, where red and blue colors indicate up- and down-regulated genes expression as measured by log2(ratio) of FPKM.

Further examination of pathways involved in β-cell function and insulin secretion revealed that many genes that were repressed by cytokines, such as *Ins1*, *Ins2*, *Pdx1*, *Isl1*, *Neurod1*, *Pax6*, were restored by I-BET treatment, as shown by the heatmap in **Figure 2E**. This unbiased analysis provided validation of gene expression results (Figure 1A-E). Interestingly, many genes in the insulin secretory pathway, including *Ins1, Ins2, Pdx1, Syntaxin, SNAP25, and Creb* all critical to GSIS, were also found to be upregulated in cells exposed to I-BET in presence of cytokine as compared to cytokines alone (CCIB vs. CCVC) (**Supplementary Figure 4**). These augmented gene expression profiles likely underlie the enhanced GSIS function in INS1 cells with I-BET treatment in the presence of cytokine exposure

### I-BET effect is mediated by antagonizing NF-kB signaling pathway

To determine the protective mechanisms of I-BET against protecting cytokine-induced β-cell dysfunction, we analyzed the differentially expressed genes induced by I-BET in cytokine-exposed cells (CCIB vs. CCVC). Gene Set Enrichment Analysis (GSEA) revealed NF-kB signaling pathway was enriched by I-BET (**Figure 3A**). Validation by RT-qPCR revealed that the downstream targets of NF-κB - *Myc* (27) and *Xiap* (28) that were suppressed by cytokines were reversed by I-BET (**Figure 3B-C**). p65 (RelA), the subunit of NF-κB transcription factor, translocates to the nucleus upon cytokine-stimulated phosphorylation and binds to the promoters/enhancers to induce expression of cytokine target genes. Hence, we tested this mechanism by examining the phosphorylation of p65 and found that it was markedly decreased by I-BET in CC-treated INS1 cells (**Figure 3D-E**). Furthermore, upstream of p65 – activation and phosphorylation of IKKA/B leads to phosphorylation and degradation of IκBα, relieving the inhibition on p65 and allowing it to translocate into the nucleus (29). We found that I-BET treatment also reduced the phosphorylation of IKK and IκBα in cytokine-exposed INS1 cells (**Figure 3F-J**), indicating an overall reduction of cytokine-induced NF-kB pathway activation. It is worth noting, however, that I-BET did not completely abolish cytokine-induced NF-kB activation, as phospho-p65 remains elevated as compared to the level observed in unstimulated INS1 cells.

**Figure 3.**
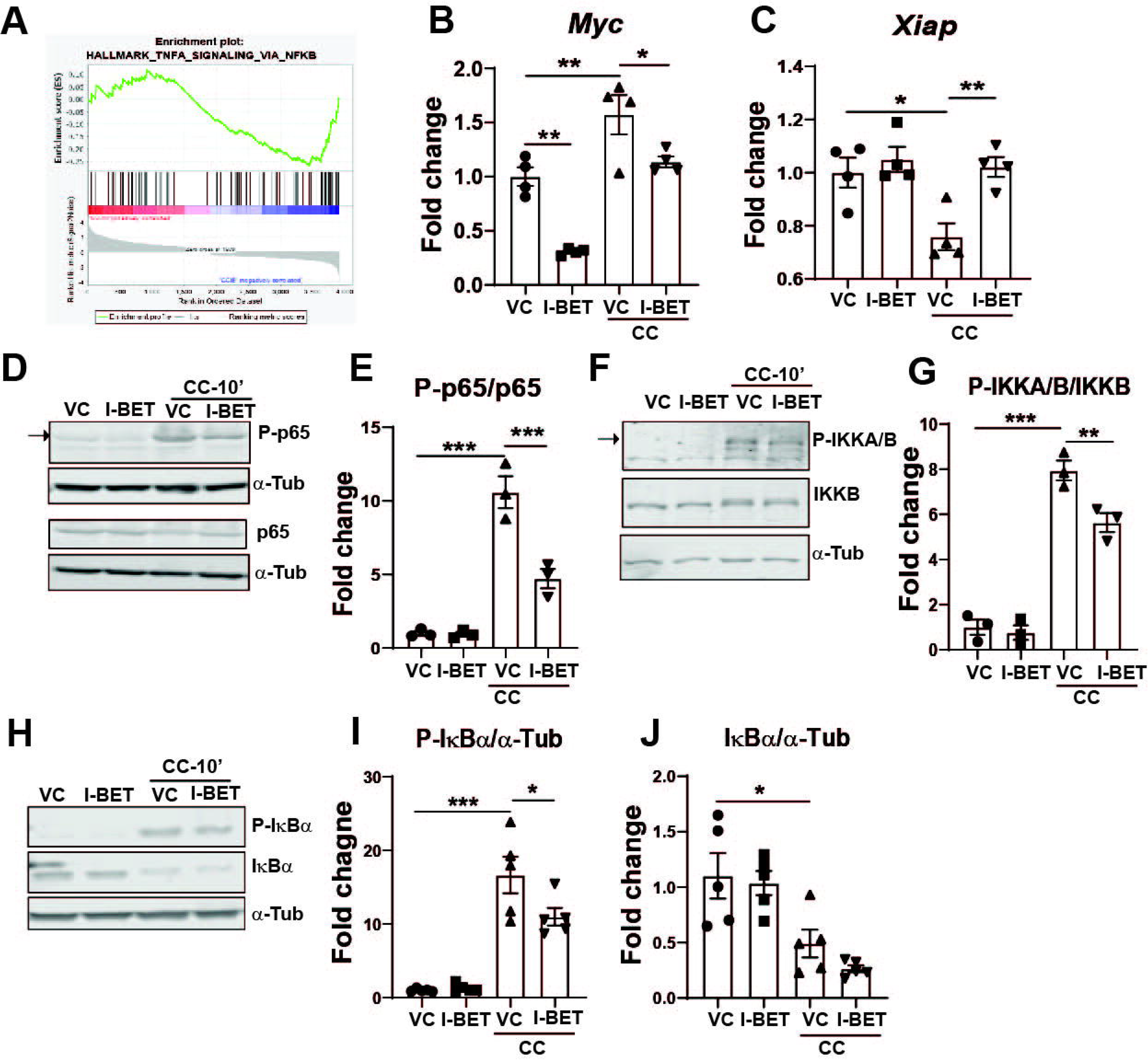
I-BET mediates its effect by antagonizing NF-kB pathway activation. **(A)** The GSEA of genes that were differentially regulated among CCIB vs CCVC, indicated TNF-α signaling via NF-kB as one of the enriched pathways represented here by enrichment plot. **(B-C)** Expression of NF-kB target genes – Myc and Xiap by RT-qPCR shown after normalization to housekeeping gene as a fold change over VC. (**D-J**) The phospho-protein and protein level by western blotting and the quantification by densitometry of 3-4 independent experiments is shown for p65 (D, E), IKKA/B (F, G) and IκBα (H-J). α-Tubulin was used as a loading control. Quantitation from densitometry is represented as a fold change over VC. The data is represented as mean ± sem (n=3-4), with at least three independent experiments. Statistical significance was calculated using one-way Anova; ***p<0.001, **p<0.01, *p<0.05.

### I-BET protects against cytokine-induced apoptosis in INS1 cells

Another finding from the GSEA of CCIB vs CCVC, was the enrichment of the apoptotic pathway (**Figure 4A**). Based on this, we tested whether I-BET can prevent β-cell apoptosis using flow cytometry analysis of Annexin-PI staining of INS1 cells. We found that cytokine exposure of 24h, but not 8h (**Supplementary Figure 5)**, was sufficient to observe significantly increased apoptosis in cytokine treated cells (CC+VC) as compared to vehicle treated controls (VC), with ~15% cells showing early apoptosis (Annexin V^+^, PI^−^), ~12.5% with late apoptosis (Annexin V^+^, PI^+^) in CC-VC cells and ~4% were dead cells (PI^+^) (**Figure 4B-C**). Importantly, I-BET significantly reduced the percentage of early (Annexin V^+^, PI^−^) and late apoptotic cells (Annexin V^+^, PI^+^) even after cytokine exposure in CC+I-BET as compared to CC+VC, without a significant change in PI^+^ cells, indicating I-BET protects against the cytokine induced β-cell apoptosis. Consistent with this, I-BET decreased the cleaved caspase-3 levels after cytokine exposure (**Figure 4D-E**), as revealed by its quantification using western blot.

**Figure 4.**
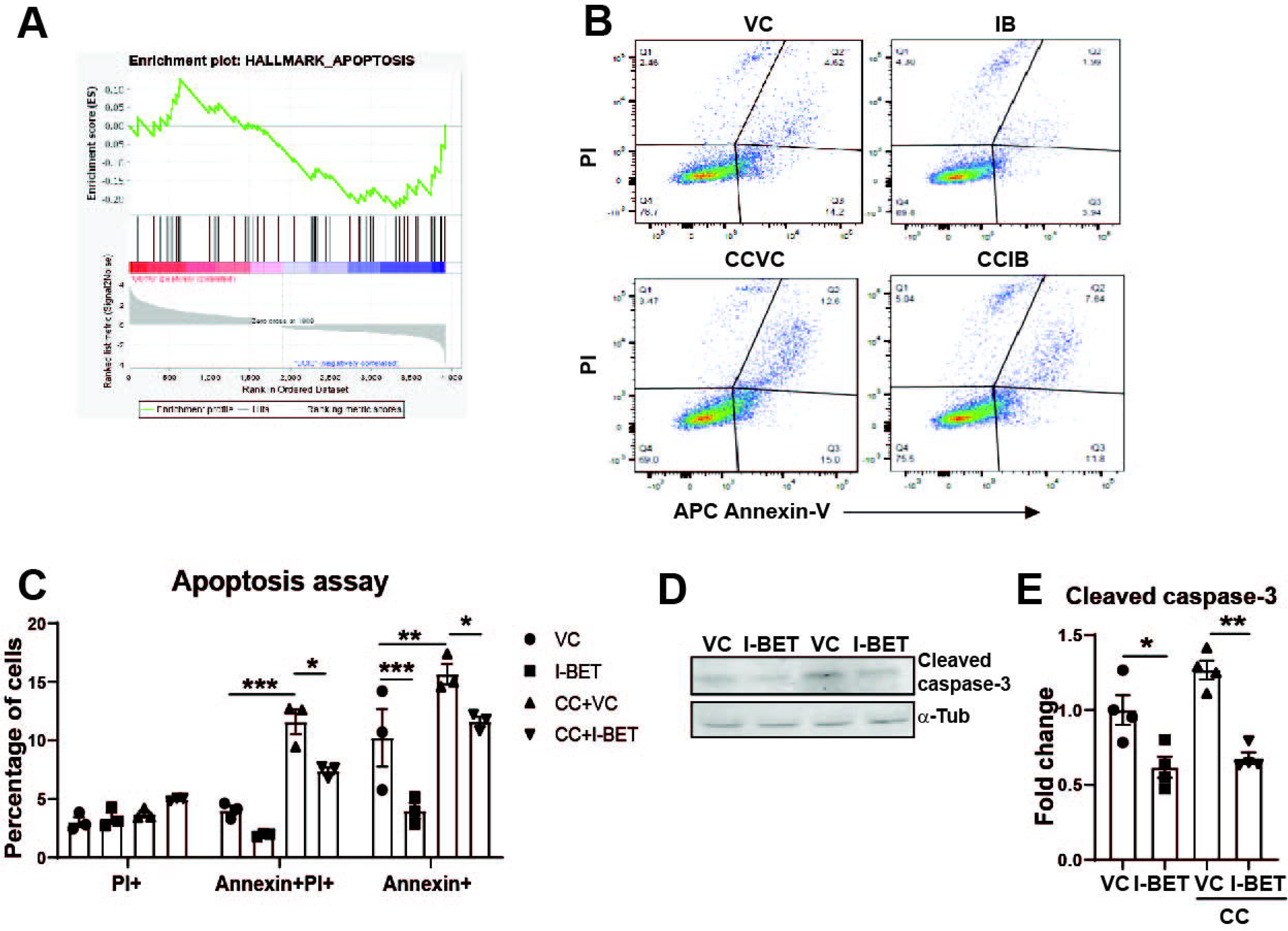
I-BET protects against cytokine-induced β-cell apoptosis in vitro. **(A)** Enrichment plot for apoptosis pathway related genes that were differentially regulated between CCIB vs CCVC by GSEA. **(B-E)** INS-1 cells pretreated with I-BET or VC (vehicle control) for 48h and then exposed to cytokine cocktail for another 24h were evaluated by PI and AnnexinV staining by flow cytometry with appropriate controls. Representative dot plot is shown in (B) for the 4 groups and quantitation of early (Annexin+) and late (Annexin+ PI+) apoptotic cells from three independent experiments is shown in (C). **(D-E)** The protein level by western blotting and the quantification by densitometry of 4 independent experiments is shown for cleaved caspase-3 and represented as a fold change over VC. α-Tubulin was used as a loading control. The data is represented as mean ± sem (n=3-4) with at least three independent experiments. Statistical significance was calculated using two-way (B) or one-way Anova (D); ***p<0.001, **p<0.01, *p<0.05.

### I-BET protects against cytokine-induced apoptosis in the multiple low-dose STZ diabetes mice model

The above experiments demonstrated that I-BET reduced the cytokine-induced apoptosis of pancreatic β-cells in vitro. We further determined if this protection translated to an in vivo effect using a well-established T1D model by multiple low-dose STZ injection (30). I-BET or vehicle (VC) was administered (30mg/Kg by gavage once daily) for 4 weeks to mice that received low dose daily STZ for 5 times at 50mg/kg (**Figure 5A**). The STZ-injected group displayed hyperglycemia starting at day 4 (data not shown), supporting the effectiveness of our diabetes model. Weekly fasting blood glucose levels were measured to assess the effect of I-BET treatment. There was a striking improvement in the fasting hyperglycemia in I-BET treated diabetic mice (**Figure 5B**). Body weight and fasting insulin levels of I-BET treated mice (STZ+I-BET) did not differ with the STZ group (STZ+VC), which were lower as compared to vehicle control (VC) as expected (**Supplementary Figure 6A-B**). Glucose tolerance test revealed significant improvement at the 30min time point between the STZ+IBET as compared to STZ+VC (**Supplementary Figure 6C**), though the insulin levels did not differ between these groups (**Supplementary Figure 6D)** suggesting that the GSIS at this late stage did not recover with I-BET treatment in vivo. Interestingly, apoptosis of pancreatic β-cell induced by STZ was significantly decreased with I-BET administration (**Figure 5C** and **5D**), consistent with the effects seen in INS1 cells presented above. Furthermore, I-BET prevented the reduction in β-cell mass (**Figure 5E**) induced by STZ. Consistent with this finding, the number of islets per imaging field in STZ+I-BET group was significantly higher than that of the STZ+VC group (**Figure 5F**), although no change in islet size distribution was observed (**Figure 5G**). This indicated that I-BET, though not effective in restoring insulin secretion, was very effective in preventing inflammation-induced β-cell apoptosis and preserving β-cell mass in the STZ-induced model of T1D.

**Figure 5.**
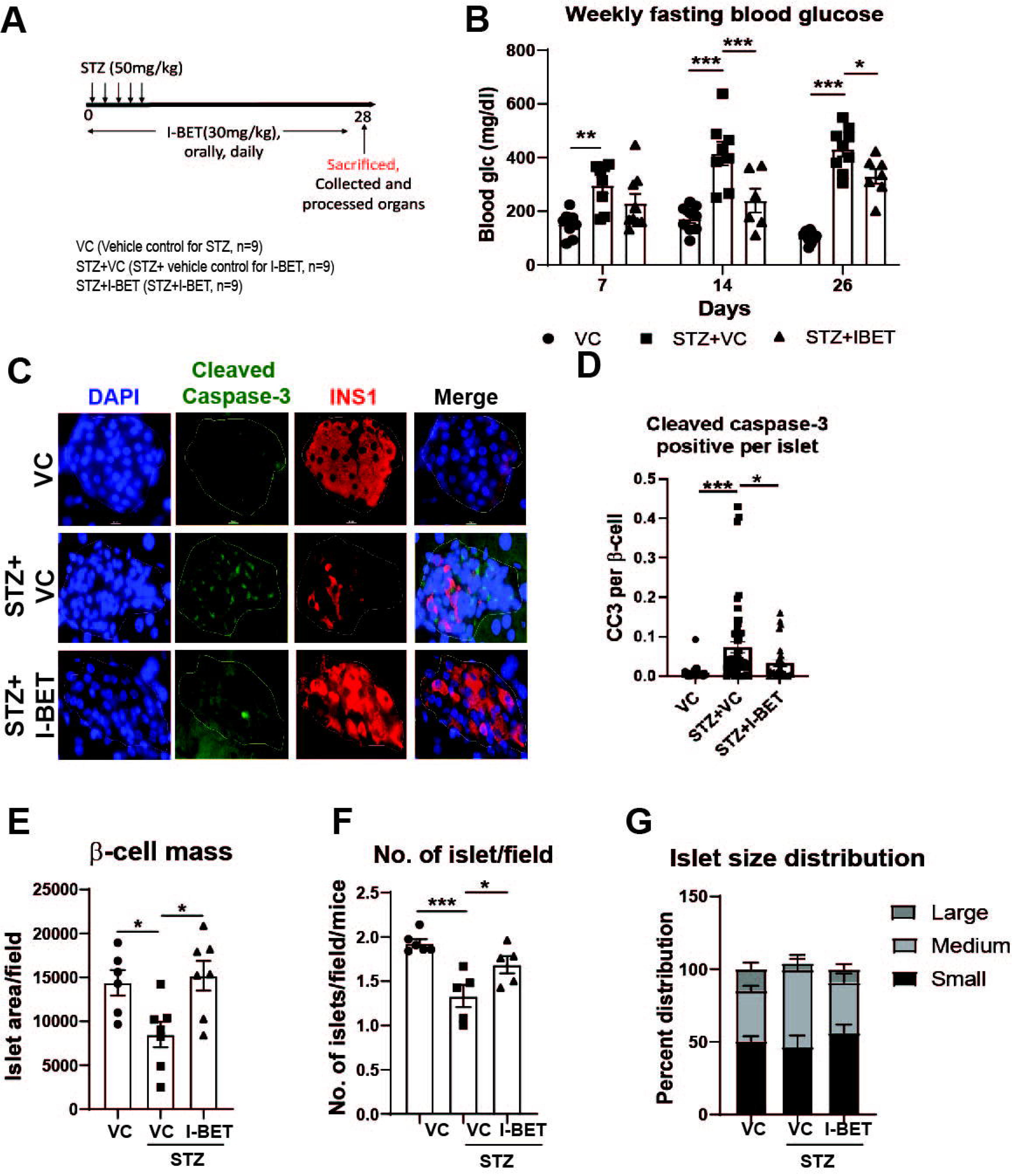
I-BET protects against cytokine-induced β-cell apoptosis in vivo. **(A)** Experimental scheme using the multiple low dose STZ mouse model. **(B)** Fasting blood glucose level (n=9 mice/group). **(C-D)** Representative sections (C) from the pancreas stained for cleaved capspase-3 (green), insulin (red), DAPI (blue) are shown. Scale bar is 10μm. Quantitation of cleaved caspase-3 signal from insulin positive β-cells in each islet is shown (D) with n=4 mice/group with 1-2 slides for each mouse. Similarly, (**E**) β-cell area, (**F**) number of islets were also quantitated in the indicated groups. (**G**) Islet size distribution was determined (n=4 mice/group with 1-2 slides for each mice) as - small, medium, large denoting the arbitrary area assigned to each islet with small being <5000, medium is 5000-30,000 and large >30,000 as quantitated in ImageJ. The data is represented as mean ± sem. Statistical significance was calculated using two-way Anova; ***p<0.001, **p<0.01, *p<0.05.

### I-BET prevents cytokine induced β-cell dysfunction in human islets

To assess if I-BET also had protective effects in human islets, non-diabetic human islets were treated with I-BET for 48h with the addition of cytokine cocktail for 24h. I-BET prevented the cytokine mediated reduction in insulin secretion (**Figure 6A**), consistent with the in vitro data from INS-1 cells. Furthermore, I-BET was also able to rescue *PDX1* and *MAFA* gene expression in cytokine treated human islets (**Figure 6B-C**) as determined by RT-qPCR. These observed protective effects of I-BET in human β-cells against cytokine-induced damage suggests its potential clinical therapeutic impact in T1D.

**Figure 6.**
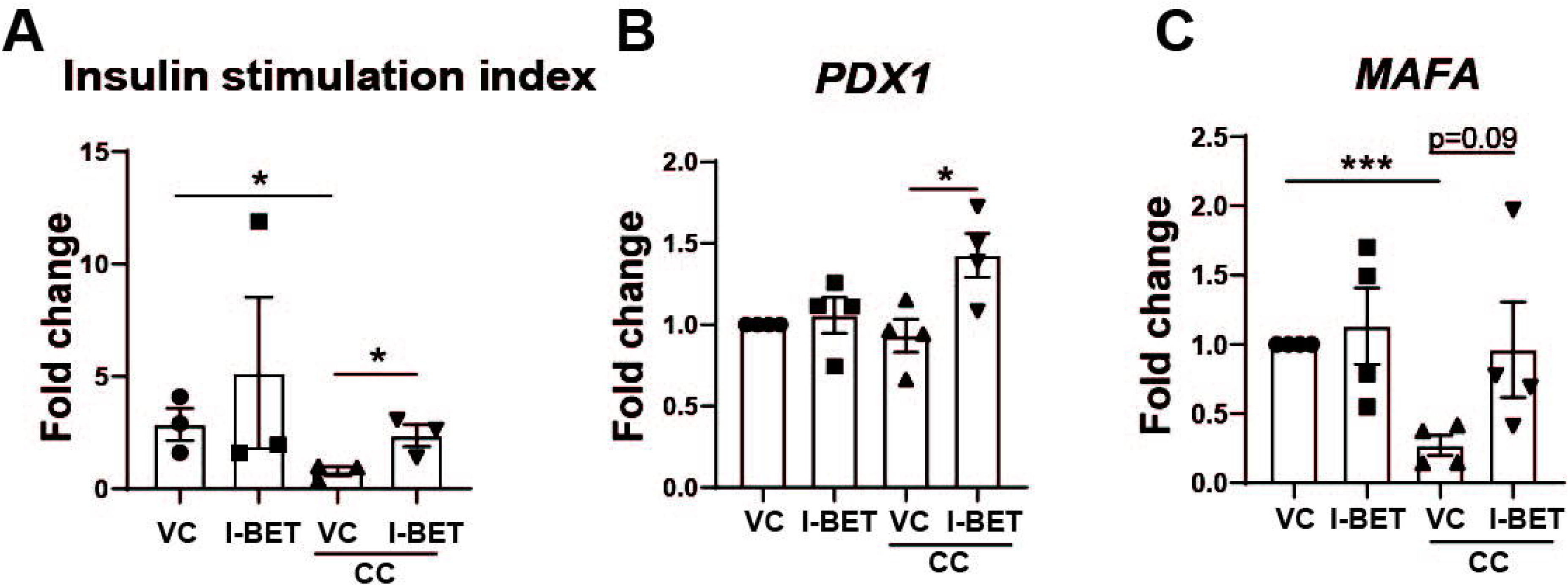
I-BET prevents cytokine induced β-cell dysfunction in human islets. **(A)** The glucose stimulated insulin secretion index expressed as a fold change over respective basal from four donor human-diabetic man islets (n=4). **(B-C)** Gene expression by RT-qPCR of (**B**) *PDX1* and (**C**) *PDX1* are shown after normalization to housekeeping gene as a fold change over VC. (n=4). The data is represented as mean ± sem. Statistical significance was calculated using Student’s t-test; *p<0.05.

## DISCUSSION

Inflammation and cytokine-induced β-cell dysfunction and apoptosis are a central mechanism in the pathogenesis of T1D (1-3). However, no targeted therapies are available in clinical practice that protect β-cells from cytokine-induced damage. In this study, we demonstrate that I-BET, a BET inhibitor has a specific effect on β-cells by antagonizing the NF-κB pathway, protecting against cytokine-induced dysfunction and cell death both in vitro, in vivo, and in human islets (**Figure 7**).

**Figure 7.**
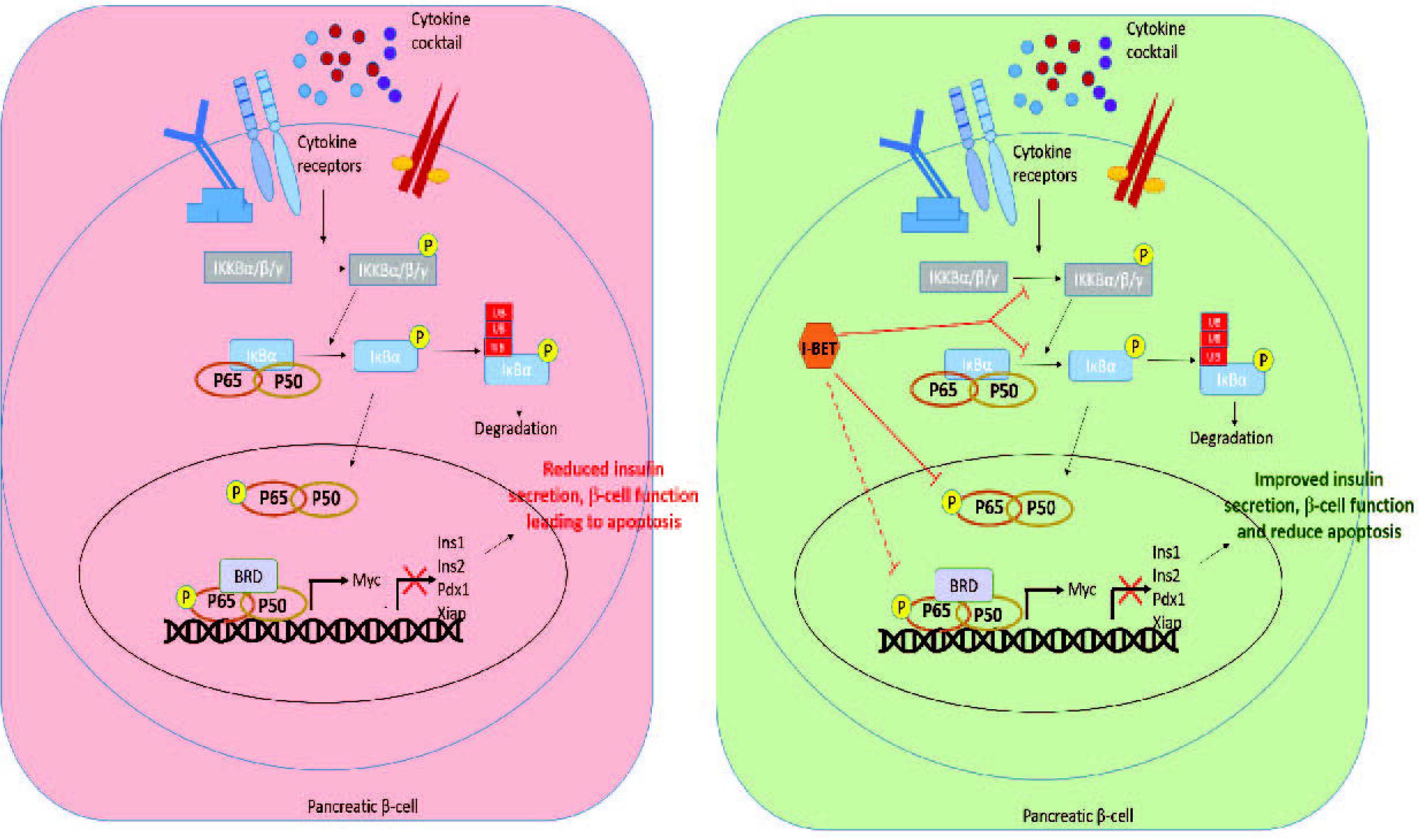
I-BET protects against cytokine-induced apoptosis by antagonizing NF-kB signaling. Under the condition of inflammation, cytokines bind to their receptors leading to phosphorylation of IKKA/B which in turn leads to phosphorylation and degradation of IκBα relieving p65 inhibition. Phospho-p65 then translocate to the nucleus, interacting with BRD proteins and regulates NF-κB target gene transcription including repression of β-cell function related genes such as *Ins1*, *Ins2* and *Pdx1*, anti-apoptotic genes –*Xiap* and activation of inflammatory genes such as *Myc*, leading to reduced insulin secretion, β-cell dysfunction and ultimately apoptosis. However, when treated with I-BET, the phosphorylation of IKKA/B, IκBα and p65 is reduced, inhibits the binding of BRD to p65, indicating an overall decrease in NF-κB pathway activity, reversing the gene expression alterations and improvement in β-cell function and reduces apoptosis. Dotted line represent data previously known, not shown in this manuscript.

Cytokine exposure leads to significant β-cell dysfunction and is often accompanied by repression of genes critical for insulin secretion, including *Pdx1* and *MafA* (31). Here we show that BET inhibition improved β-cell function by increasing the expression of *Pdx1*, *MafA*, and *Pax6* involved in maintaining β-cell secretory function. Interestingly, there were others such as *Nkx2.2* and *Nkx6.1* that were not responsive to BET inhibition in the presence of cytokines, indicating that bromodomain dependent regulation in β-cells is likely gene-specific and that there is likely a differential susceptibility of these genes to cytokine induced suppression. Importantly, similar beneficial results of improved GSIS and an increase in expression of *PDX1* and *MAFA* in I-BET treated human islets exposed to cytokines attests to the translational significance of BET inhibition in the context of T1D.

NF-κB signaling is known to be the major pathway responsible for β-cell apoptosis (32; 33). Many studies in other tissues undergoing inflammation have demonstrated activation of NF-κB signaling that is regulated by bromodomain proteins and display a beneficial response to I-BET treatment (34–37), though to date none have been demonstrated in β-cells. Brd4, a target of I-BET is known to co-activate NF-κB signaling by interacting with acetylated RelA (p65), increasing its transactivation activity and stability in cancer cells (35; 38). Transcriptomic analysis of islets from NOD mice treated with I-BET showed a differential expression of NF-κB targets (14) suggesting at least a subset of NF-κB target genes in β-cells are also regulated by BET proteins. Herein, we identify a potential mechanism of I-BET action in protecting β-cells from cytokine induced dysfunction and apoptosis wherein the critical phosphorylation of p65 (RelA), required for NF-κB translocation and transactivation, is decreased by I-BET treatment via a reduction in phosphorylation of IKKA/B and IκB. This is consistent with results of BET inhibition in other cell types in lymphomas (B cells), rheumatoid arthritis (synoviocytes), and cancers (microglia) (39–41). In addition to the regulation of phospho-p65 (RelA), there are other potential mechanisms whereby I-BET treatment can downregulate NF-κB signaling. For instance, BET inhibitors in a dose-dependent manner inhibit the binding of Brd4 to acetylated RelA, a critical step in the formation of RelA:P-TEFb transcription-activation complex (38; 42). Another potential mechanism whereby I-BET could repress NF-κB signaling is by preventing the formation of RelA-dependent super-enhancers that are critical in activation of the cytokine-inflammatory pathway target genes (43), the identification and regulation of which will be the subject of future studies.

Despite β-cell dysfunction with 8h of cytokine-exposure, an increase in apoptosis required a longer exposure of 24h, indicating apoptosis as a late event. However, at the 24h time point, with the onset of cytotoxicity and apoptosis, not surprisingly, I-BET was unable to rescue cytokines-induced β-cell dysfunction including insulin secretion and β-cell related gene expression. This inability of I-BET to improve β-cell function at later time points could be underpinning the modest glycemic response and an inability to rescue the loss of insulin secretion during the GTT in our STZ-diabetic mice. Importantly, similar to the in vitro studies, I-BET was successful in preventing apoptosis and preservation of β-cell mass in the STZ-diabetes mice, which could have significant translational value in early human T1D management. A limitation with the systemic administration of I-BET to STZ-induced diabetic mice is the likely role of BET inhibition in infiltrating immune cells in the islets, and the role of other tissues, such as the liver, in ameliorating hyperglycemia, making it difficult to distinguish the cell-autonomous effects of I-BET in β-cells. Though the studies in INS1 cells strongly support a cell-autonomous effects of I-BET in β-cell in vivo, similar experiments in mice with genetic deletion of specific BET proteins or the design of specific inhibitors to the individual BET proteins will resolve this question.

Global transcriptomic analysis revealed that I-BET was able to prevent many but not all transcriptomic changes induced by cytokine exposure. This could be a result of the differential dependence of cytokine-induced transcriptional changes on BET-dependent and BET-independent mechanisms and on relative roles of Brd2 and Brd4, arguing that a combination of agents may be needed to fully repress the deleterious effects of cytokines. While the current study was not designed to determine the relative role of the various BET proteins in the protective effect of I-BETs, a more specific inhibitor that can differently affect BET proteins could likely be more efficacious.

Although the translational benefits of BET protein inhibition in oncologic and inflammatory diseases are being assessed in ongoing clinical trials, its role in human diabetes awaits more translational work. The results from our study, especially the protective effects of preserving β-cell function in cytokine-exposed human islets is encouraging and should trigger additional mechanistic and translational studies to explore the role of BET protein inhibition as potential therapy in T1D.

## Guarantor

V.K. Yechoor is the guarantor of this work and, as such, had full access to all the data in the study and takes responsibility for the integrity of the data and the accuracy of the data analysis.

## Author contributions

VN, JL, RL, MG, FL, RJ, PY, RB, KM, MM, VY: acquisition of the data or the analysis and interpretation of information. VN, MM, VY: involvement in the conception, hypotheses delineation, and design of the study. VN, KM, VY: writing the article or substantial involvement in its revision prior to submission.

## Prior presentation information

Parts of this study were presented in abstract form at the 80th Scientific Sessions of the American Diabetes Association (https://doi.org/10.2337/db20-2036-P). A non–peer-reviewed version of this article was posted on the bioRxiv preprint server (link) on 4 November 2020.

