## Supplementary material for "Bromodomain protein inhibition protects β-cells from cytokine-induced death and dysfunction via antagonism of NF-κB pathway": Online Supplementary material

^1^Diabetes and Beta Cell Biology Center, Division of Endocrinology and Metabolism, ^2^Division of Cardiology, Department of Pediatrics, Children’s Hospital of Pittsburgh, University of Pittsburgh, Pittsburgh, PA. ^3^Institute of Cellular Therapeutics, Allegheny Health Network, 320 East North Avenue, Pittsburgh, PA, ^4^Department of Diabetes Complications and Metabolism, Diabetes and Metabolism Research Institute, City of Hope, CA. *Corresponding

**Short running title**: I-BET protects against β-cell apoptosis

**Corresponding author**: Dr. Vijay Yechoor, MD

Professor of Medicine

Division of Endocrinology, Diabetes and Metabolism.

Director of Diabetes and Beta Cell Biology Center

Endowed Chair for Diabetes and Metabolic Bone Disease

University of Pittsburgh

Pittsburgh, PA 15213

412-383-4251

**ONLINE SUPPLEMENT**

**Supplementary Figure 1. Gene expression changes with I-BET at 8h of cytokine exposure.**

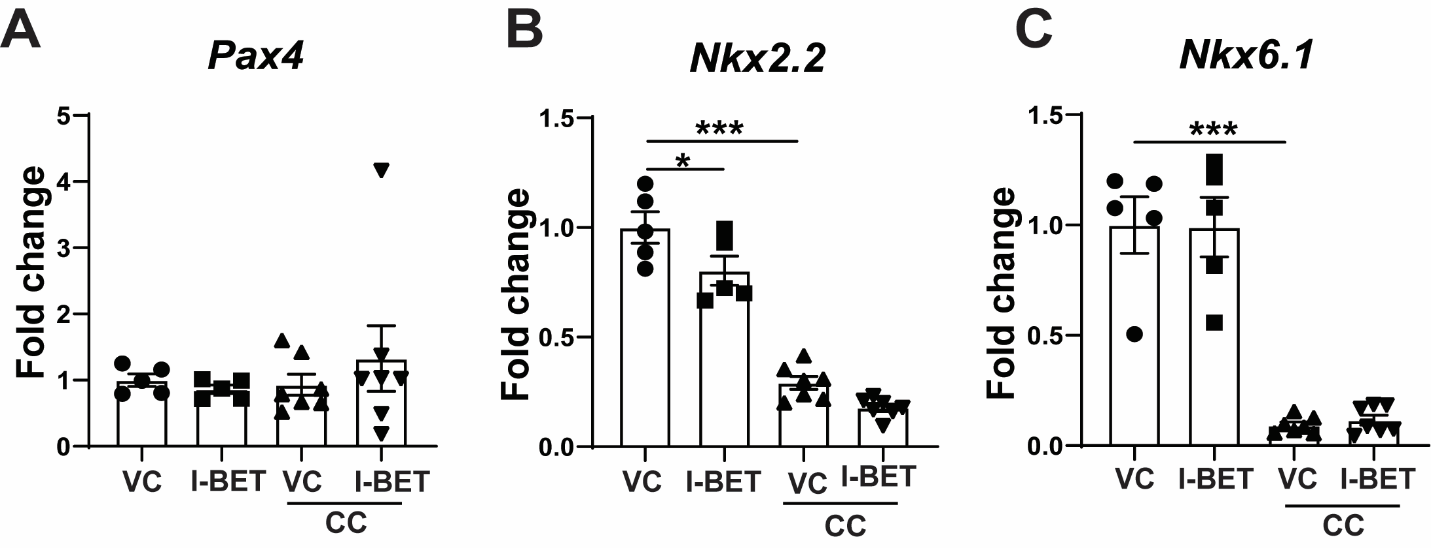

**(A-C)** Gene expression by RT-qPCR of (**A**) *Pax4*, (**B**) Nkx2.2 (**C**) Nkx6.1 are shown after normalization to housekeeping gene as a fold change over VC (vehicle control). The data is represented as mean ± sem (n=5-6), with at least three independent experiments. Statistical significance was calculated using one-way Anova; ***p<0.001, **p<0.01, *p<0.05.

**Supplementary Figure 2. I-BET does not rescue cytokine-induced decrease in β-cell function at 24h time point.**

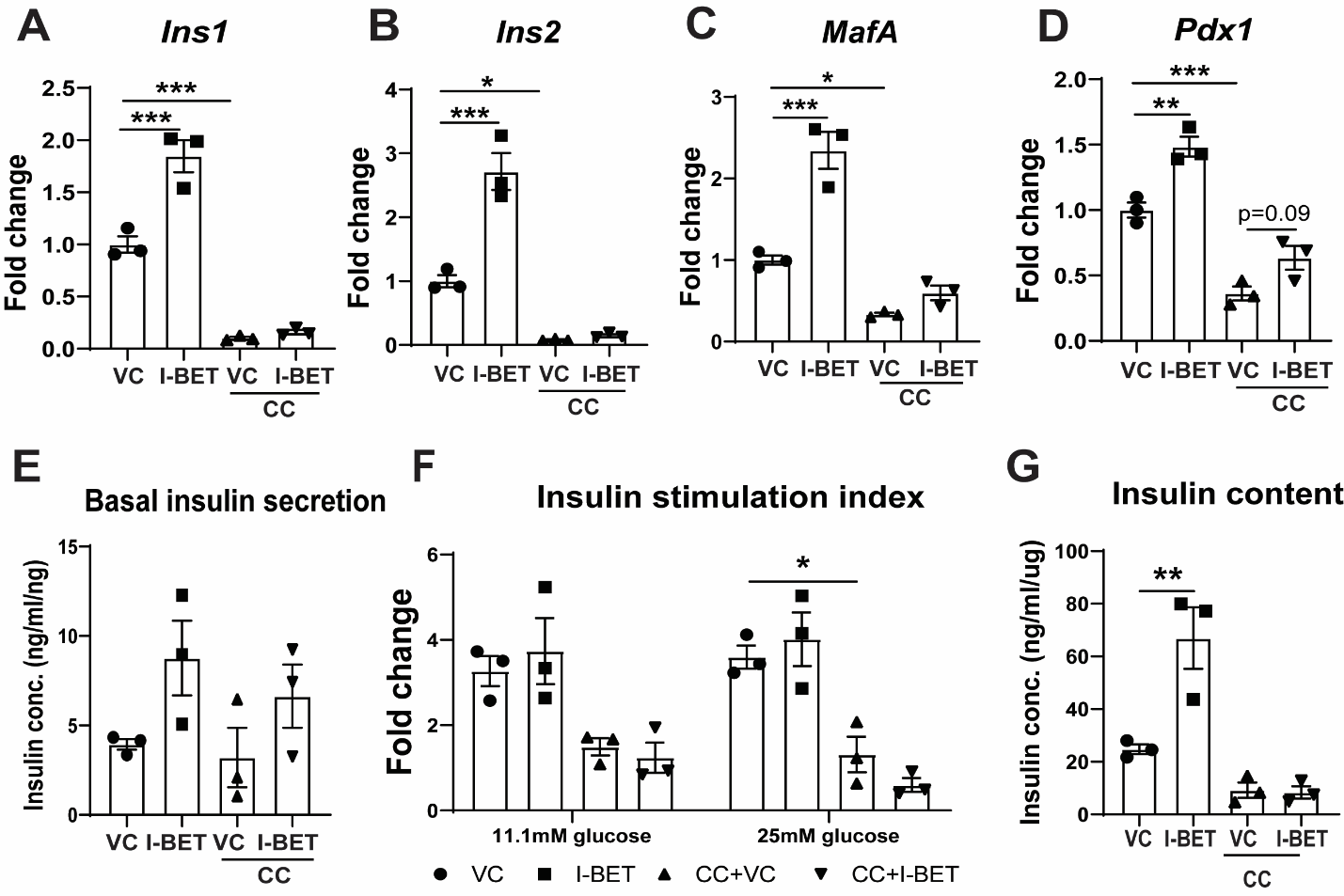

**(A-D)** Gene expression by RT-qPCR of (**A**) *Ins1*, (**B**) *Ins2*, (**C**) *MafA*, (**D**) *Pdx1*, are shown after normalization to housekeeping gene as a fold change over VC. (**E-G**) The secreted insulin from INS1 cells in basal 2.8mM glucose (E) and after incubation in indicated glucose concentrations (F) is shown. Insulin secretion is represented as insulin stimulation index (F) a fold change over the respective levels from the basal 2.8mM glucose. (**G**) Insulin content measured in INS1 cell lysates normalized to cellular DNA. The data is represented as mean ± sem (n=3), with at least three independent experiments. Statistical significance was calculated using one-way Anova; ***p<0.001, **p<0.01, *p<0.05.

**Supplementary Figure 3. Pathways altered by I-BET in presence and absence of cytokines.**

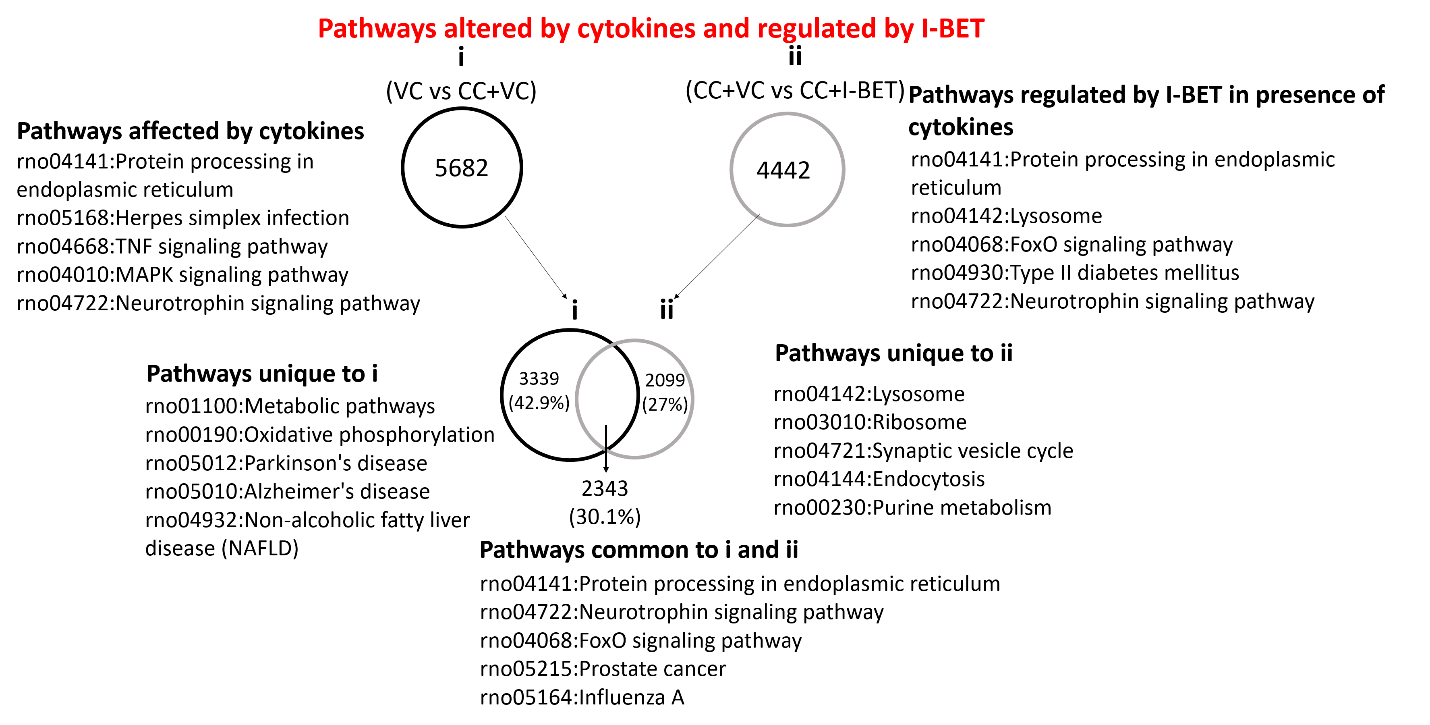

To determine pathways altered by both cytokines (VC vs CC+VC) and I-BET (CC+I-BET), the respective differential expression gene lists were plotted as Venn diagram. Briefly, 5682 genes were differentially expressed among VC and CC+VC and 4443 among CC+VC and CC+I-BET. Among these 30.1% genes were regulated by both CC and I-BET where as 42.9% were unique to A and 27% to B.

**Supplementary Figure 4. I-BET treatment regulates insulin secretion pathway in cytokine-exposed INS1 cells.**

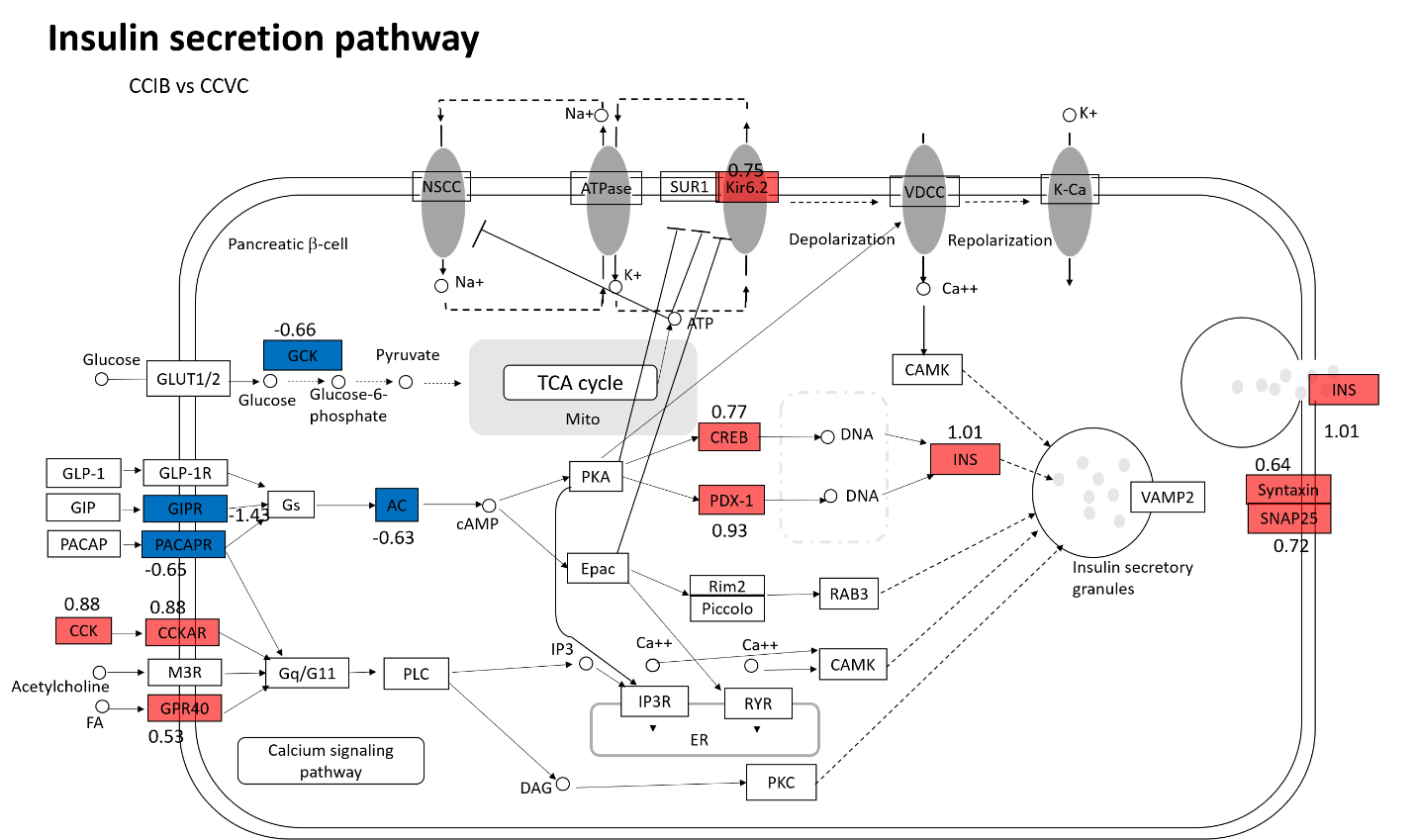

Insulin secretion pathway was enriched in the differentially expressed genes by CCIB. Red and blue represent up-and down-regulated genes in comparison of CCIB vs CCVC. The log2 fold change is shown below the gene.

**Supplementary Figure 5. 8h of cytokine exposure is insufficient for apoptosis induction.**

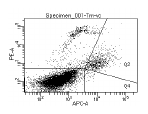

6%

7.7%

0.6%

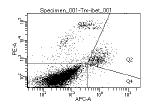

5.1%

6.1%

0.5%

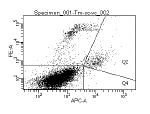

7%

4%

0.4%

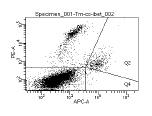

9.8%

3.9%

0.5%

**VC**

**I-BET**

**CC+VC**

**CC+ I-BET**

**B**

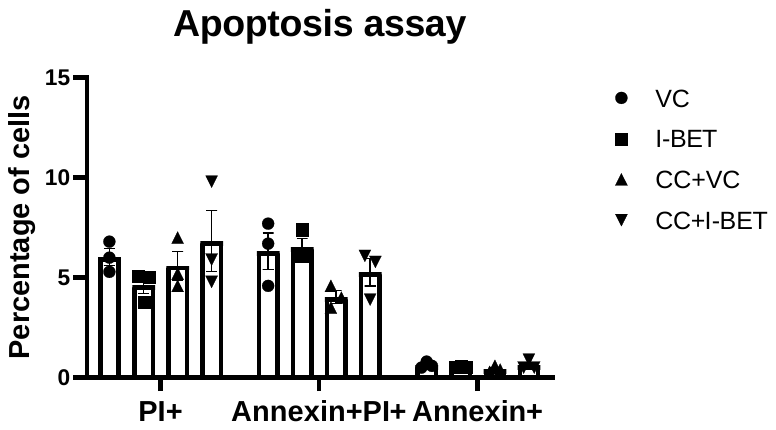

**A**

**APC Annexin-V**

**PI**

INS-1 cells pretreated with I-BET or VC (vehicle control) for 48h and then exposed to cytokine cocktail for another 8h were evaluated by PI and AnnexinV staining by flow cytometry with appropriate controls. Representative dot plot is shown in **(A)** for the 4 groups and quantitation of early (Annexin+) and late (Annexin+ PI+) apoptotic cells from three independent experiments is shown in **(B)**. The data is represented as mean ± sem (n=3) with at least three independent experiments. Statistical significance was calculated using two-way (B) or one-way Anova (D); ***p<0.001, **p<0.01, *p<0.05.

**Supplementary Figure 6. Effect of I-BET on multiple low dose STZ mouse model of T1D.**

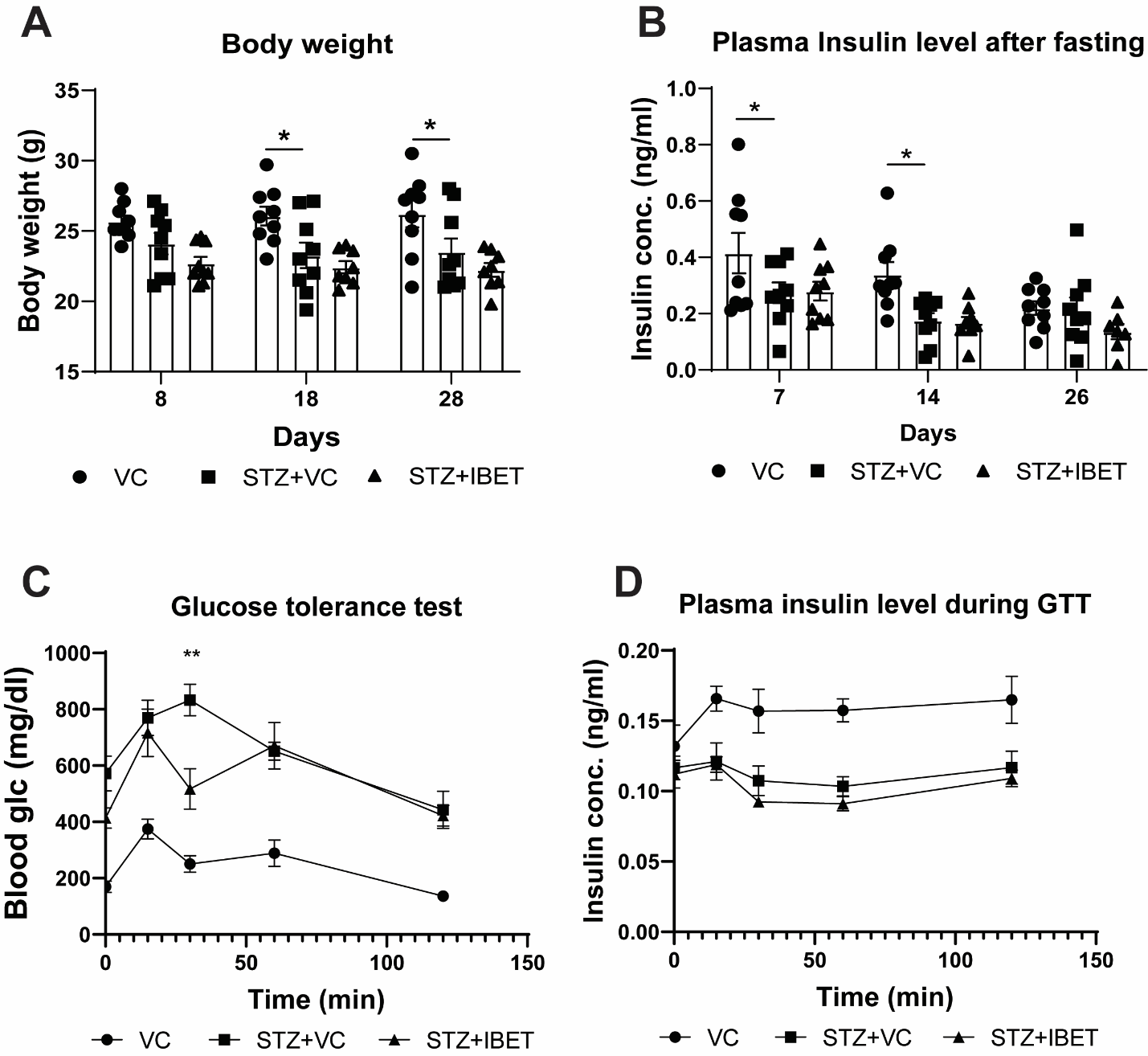

(**A)** Body weight **(B)** Plasma insulin level in fasted mice in indicated groups. **(C-D)** Glucose tolerance test (GTT) in fasted mice with glucose (C) and insulin (D) levels at indicated time points. The data is represented as mean ± sem (n=9). Statistical significance was calculated using two-way Anova; ***p<0.001, **p<0.01, *p<0.05 for STZ+VC and STZ+I-BET group and #<0.05 for VC and STZ+VC group.

**Supplementary Table 1. List of Primers used.**

| **S.No.** | **Gene name** | **Primer name** | **Sequence** |
| --- | --- | --- | --- |
| 1. | Insulin 1 | rt-Ins1-FP | ACAGCACCTTTGTGGTCCTC |
| 2. |  | rt-Ins1-RP | GTGCAGCACTGATCCACAAT |
| 3. | Insulin 1 | rt-Ins2-FP | TGTGGTTCTCACTTGGTGGA |
| 4. |  | rt-Ins2-RP | ATGCTGGTGCAGCACTGA |
| 5. | Pancreatic and Duodenal Homeobox 1 | rt-Pdx1-FP | AGCTCACGCGTGGAAAAG |
| 6. |  | rt-Pdx1-RP | GTACGGGTCCTCTTATTCTCCTC |
| 7. |  | hu-Pdx1-FP | GGAAAACCCGCTCTCTCAGG |
| 8. |  | hu-Pdx-1-RP | CCAAGGTGGAGTGCTGTAGG |
| 9. | MAF BZIP Transcription Factor A | rt-Mafa-FP | AGCAAGGAGGAGGTCATC |
| 10. |  | rt-Mafa-RP | CGTATTTCTCCTTGTACAGG |
| 11. |  | hu-Mafa-FP | AGAGCGAGAAGTGCCAACTC |
| 12. |  | hu-Mafa-RP | TTGTACAGGTCCCGCTCTTT |
| 13. | Paired Box 6 | rt-Pax6-FP | CCAGTTTTCAGAGCCACGTAT |
| 14. |  | rt-Pax6-RP | ACTCCGCTGTGACTGTTCTG |
| 15. | NK6 Homeobox 1 | rt-Nkx6.1-FP | ATGGGAAGAGAAAACACACCAGAC |
| 16. |  | rt-Nkx6.1-RP | TAATCGTCGTCGTCCTCCTCGTTC |
| 17. | NK2 Homeobox 2 | rt-Nkx2.2-FP | CAGCGACAACCCCTACACTC |
| 18. |  | rt-Nkx2.2-RP | GCTTTGGAGAAGAGCACTCG |
| 19. | Paired Box 4 | rt-Pax4-FP | AGATGTTCCAGTGACACCACA |
| 20. |  | rt-Pax4-RP | CACAGGAAGGAGGGAGTGG |
| 21. | Myc | Rt-Myc-FP | GCTCCTCGCGTTATTTGAAG |
| 22. |  | Rt-Myc-RP | GCATCGTCGTGACTGTCG |
| 23. | Xiap | Rt-Xiap-FP | GCTTGCAAGAGCTGGATTTT |
| 24. |  | Rt-Xiap-RP | TGGCTTCCAATCCGTGAG |

**Supplementary Table 2. Human islet donor details**

| **S.No.** | **Unique Identifier number** | **Donor Age** | **Donor Sex** | **Donor BMI** | **History of Diabetes** | **Islet purity** |
| --- | --- | --- | --- | --- | --- | --- |
| 1. | AGH20190726 | 49 | Male | ‘obese’ | No | 90% |
| 2. | SAMN13836615 | 58 | Male | 23.2 | No | 90% |
| 3. | SAMN15314807 | 27 | Male | 25.3 | No | 85% |
| 4. | SAMN15337453 | 26 | Male | 24.2 | No | 98% |
